# CRISPR/Cas12a-mediated allele engineering of *SmAPRR2* and *SmGLK2* reveals complementary control of fruit peel and flesh chlorophyll pigmentation in eggplant

**DOI:** 10.64898/2026.07.28.741273

**Authors:** Marina Martínez-López, Andrea Solana, Andrea Arrones, Laura Toppino, Santiago Vilanova, Mariola Plazas, Jaime Prohens, Pietro Gramazio

## Abstract

Eggplant (*Solanum melongena* L.) displays extensive fruit color diversity, in which chlorophyll-related pigmentation contributes to both external appearance and market value. Previous genetic studies identified *SmAPRR2* and *SmGLK2* as major candidate genes controlling uniform green pigmentation and green netting in fruit, respectively, but their individual and combined functional contributions had not been validated through targeted mutagenesis in a common genetic background. Here, we established a multiplex CRISPR/Cas12a system in eggplant accession MEL3, representing, to our knowledge, the first application of this nuclease for genome editing in eggplant. Transformation efficiency was 2.0%, but all 15 genotyped regenerants were edited, yielding four *SmAPRR2* and six *SmGLK2* alleles. Segregation and crossing enabled the recovery of six transgene-free lines carrying single or combined edited alleles. Disruption of *SmGLK2* abolished the reticulated green netting pattern while preserving a uniformly green peel and the internal green ring. Conversely, disruption of *SmAPRR2* reduced the background uniform peel pigmentation and eliminated the green ring while retaining green netting. Double mutants carrying disruptive alleles at both loci produced white fruits lacking internal green pigmentation, whereas putatively hypomorphic *SmAPRR2* and *SmGLK2* alleles generated intermediate phenotypes. Whole-genome resequencing identified only two predicted off-target sites under a canonical TTTV PAM search allowing up to four mismatches. Both were fully covered, and no edited-line-specific candidate variants were detected. These findings establish complementary and partially separable roles for *SmAPRR2* and *SmGLK2* in fruit peel and flesh chlorophyll pigmentation and demonstrate the potential of Cas12a for functional genomics, allele engineering, and precision breeding in eggplant.

## 1. Introduction

Eggplant (*Solanum melongena* L.) is a globally important vegetable crop displaying remarkable diversity in fruit color patterns, a key determinant of market appeal and consumer preference (Page et al., 2019). Fruit coloration in eggplant presents a wide range of pigmentation patterns, predominantly influenced by the combined accumulation and spatial distribution of anthocyanins and chlorophylls. While anthocyanins contribute to the purple coloration, chlorophyll accumulation underlies green pigmentation that is visible in both the presence and absence of anthocyanins (Page and Chapman, 2021). Two major genes, the *ARABIDOPSIS PSEUDO RESPONSE REGULATOR2-LIKE* gene *(SmAPRR2)* and the *GOLDEN2-LIKE2* gene *(SmGLK2)*, have been proposed as key regulators of chlorophyll-related fruit pigmentation, affecting green background color and reticulated green netting, respectively (Arrones et al., 2022, 2024).

*SmAPRR2* encodes an APRR-like pseudo-response regulator, which is involved in pigment accumulation and chloroplast development in various plant species, including solanaceous crops (Pan et al., 2013; Jiao et al., 2017). In eggplant, *SmAPRR2* has been associated with the presence and intensity of uniform green pigmentation across the fruit peel (Arrones et al., 2022; Fang et al., 2023). In addition, its genomic proximity to a previously reported locus controlling the green ring in the flesh (Portis et al., 2014) suggests that *SmAPRR2*-dependent chlorophyll regulation may also extend to internal fruit tissues adjacent to the peel. Similar roles of *APRR2-like* genes in chlorophyll accumulation have been reported in cucurbit crops such as cucumber, melon, pumpkin, watermelon, and chayote (Jiao et al., 2017; Oren et al., 2019; Gebretsadik et al., 2024; Cheng et al., 2025).

The *SmGLK2* gene belongs to the GARP family of MYB transcription factors, which are widely conserved and have been linked to chloroplast biogenesis across various plant species, including tomato (*S. lycopersicum*), maize (*Zea mays*), rice (*Oryza sativa*), and *Arabidopsis thaliana* (Rossini et al., 2001; Powell et al., 2012; Tachibana et al., 2024; Zhang, Zhang, Zeng et al., 2024). In Solanaceae, *GLK* genes appear to have undergone an early duplication that gave rise to two orthologous lineages, *GLK1* and *GLK2*, followed by partial functional specialization: *GLK1* generally shows stronger expression in vegetative tissues, whereas *GLK2* tends to predominate in fruits (Brand et al., 2014; Nguyen et al., 2014). Consistent with this functional divergence, expression data in the eggplant wild ancestor *S. insanum* showed that both genes are expressed at similar levels in seedlings, whereas *GLK2* expression is higher than *GLK1* in young leaves, flowers, and especially fruits, where *GLK1* expression is nearly undetectable (Arrones et al., 2024). In eggplant, *SmGLK2* has been associated with the green netting phenotype of the fruit peel, which is characterized by an irregular, non-uniform distribution of green pigmentation forming a reticulated pattern, particularly pronounced in the proximal region of the fruit (Arrones et al., 2024). In tomato, the orthologous *SlGLK2* gene underlies the Uniform Ripening (*U*) locus, and mutations at this locus are associated with the loss of green shoulder pigmentation, a phenotype that shows parallels with the green netting trait in eggplant (Powell et al., 2012). In both species, variation in chlorophyll-related pigmentation affects fruit appearance and ripening characteristics, thereby influencing marketability and visual quality and highlighting *SmGLK2* as a promising target for breeding programs aimed at modifying fruit color patterns (Powell et al., 2012; Arrones et al., 2024).

In our previous studies, *SmAPRR2* was identified as a strong candidate gene associated with uniform chlorophyll pigmentation in the fruit peel using a genome-wide association study (GWAS) in an eggplant multi-parent advanced generation inter-cross (MAGIC) population and sequence analysis of a diverse germplasm collection (Arrones et al., 2022). Consistently, *SmAPRR2* was independently fine-mapped as a gene regulating green/white rind color in eggplant, further supporting its central role in chlorophyll-related fruit peel pigmentation (Fang et al., 2023). Similarly, *SmGLK2* was identified as a candidate gene associated with the irregular green netting trait in eggplant fruits through GWAS in the eggplant MAGIC population, combined with additional bulk segregant analysis by sequencing (BSA-seq) in an F2 population, fine-mapping in advanced backcross populations, and expression analysis (Arrones et al., 2024). Building on this genetic framework, the present study uses targeted mutagenesis to dissect the individual and combined contributions of both genes in a common genetic background, providing direct functional evidence for their complementary roles in eggplant fruit pigmentation.

Although the first successful *Agrobacterium*-mediated transformations in eggplant were achieved over 35 years ago (Guri and Sink, 1988; Rotino and Gleddie, 1990), progress in improving transformation protocols and their efficiencies has been slow. Despite continuous efforts to optimize these methods (Magioli et al., 2000; Vinod Kumar and Rajam, 2005; Prakash et al., 2007; Sagare and Mohanty, 2012), it has only been in recent years that more reliable and effective protocols have emerged, offering a more solid foundation for genome editing workflows (Maioli et al., 2020; Khatun et al., 2022; Wang et al., 2022). However, genome editing in eggplant remains relatively underexplored. To our knowledge, so far only nine published studies have addressed this topic, all based on Cas9-mediated editing (Maioli et al., 2020; Wang et al., 2022; Kodackattumannil et al., 2023; Phad et al., 2024; Sagarbarria et al., 2023; Ferrero et al., 2024; Lee et al., 2024; Liao et al., 2024; Zhang, Zhang, Yan et al., 2024; Toppino et al., 2025), and no standardized protocol for functional validation in this species has yet been established. This contrasts with tomato, where genome editing is widely and routinely applied for gene function studies (Saini et al., 2025; Sakthivel et al., 2025).

Overall, this limited body of work highlights both the promise and the challenges still faced in applying CRISPR/Cas technology to eggplant, as transformation and editing efficiencies remain low and are often highly genotype-dependent (Sagarbarria et al., 2023). Moreover, the lack of standardized protocols for eggplant transformation has led to considerable variability in results, limiting reproducibility and the broader application of these technologies in breeding programs. In this context, the use of CRISPR/Cas12a represents a significant advancement. Cas12a offers several advantages over the widely used Cas9, including higher specificity and lower tendency for off-target effects, making it an attractive tool for genome editing in challenging species such as eggplant (Swarts and Jinek, 2018; Paul and Montoya, 2020). Although Cas12a has not previously been applied to eggplant, its distinct features, including a T-rich PAM and staggered DNA cleavage, make it a promising candidate for targeted and efficient genome editing in this crop, particularly given the persistent limitations associated with transformation efficiency and protocol standardization (Banakar et al., 2020; Paul and Montoya, 2020).

In this study, we implemented a multiplex CRISPR/Cas12a strategy to functionally validate *SmAPRR2* and *SmGLK2* in eggplant, representing, to our knowledge, the first report of Cas12a-mediated genome editing in this crop. To evaluate both genes in a common genetic background, we selected the anthocyanin-free MEL3 accession, whose fruits combine a uniform green peel background with an irregular dark green netting pattern. By generating transgene-free single- and double-edited lines in the *S. melongena* MEL3 background, we resolved the individual and combined contributions of both genes to chlorophyll-related pigmentation of the fruit peel and flesh. We also evaluated predicted off-target sites using whole-genome resequencing data from selected edited lines. Our results provide direct loss-of-function validation of both genes, reveal complementary control of green background pigmentation, green netting, and internal green ring, and support Cas12a-mediated multiplex editing as a tool for functional genomics and targeted allele diversification in eggplant.

## 2. Results

### 2.1. Generation and molecular characterization of CRISPR/Cas12a-edited lines

Following *Agrobacterium*-mediated transformation, adventitious shoot regeneration was observed in some infected explants (Figures 1A, B), and *DsRed* fluorescence allowed the visual identification of putative transgenic regenerants. In this way, fluorescence signal was detected in regenerated shoots (Figure 1C), leaves (Figure 1D), and roots (Figure 1E), confirming reporter expression in different tissues. Overall transformation efficiency was low, with transgenic regenerants obtained from six explants out of a total of 300 initially infected explants, corresponding to an efficiency of 2.0% (95% exact binomial confidence interval: 0.7-4.3%). All genotyped transgenic regenerants carried CRISPR-induced mutations, indicating that editing was successfully achieved among the recovered transformants, although this result should be interpreted cautiously given the limited number of independent events. Representative *DsRed*-positive regenerants derived from these explants were selected for genotyping, prioritizing independent regeneration events and distinct allelic profiles. Among the 15 genotyped plants, all carried CRISPR-induced mutations at *SmGLK2* (100% editing frequency), whereas mutations at *SmAPRR2* were detected in nine plants (60%) (Table S1). Based on the sequencing and the ICE analysis, three T_0_ lines (T_0__1, T_0__3, and T_0__9) were selected for self-pollination and further characterization. These lines were chosen because they captured the main editing profiles observed among the genotyped regenerants and showed high predicted knockout (KO) scores at the targeted loci. T_0__1 carried mutations only at *SmGLK2*, whereas T_0__3 and T_0__9 carried mutations at both *SmAPRR2* and *SmGLK2*. In addition, T_0__9 showed an editing profile compatible with a large deletion of approximately 918 bp in *SmGLK2*. An overview of the genealogy of the selected edited lines, including the subsequent selfing, crossing, and segregation steps, is provided in Figure S1.

**Figure 1.**
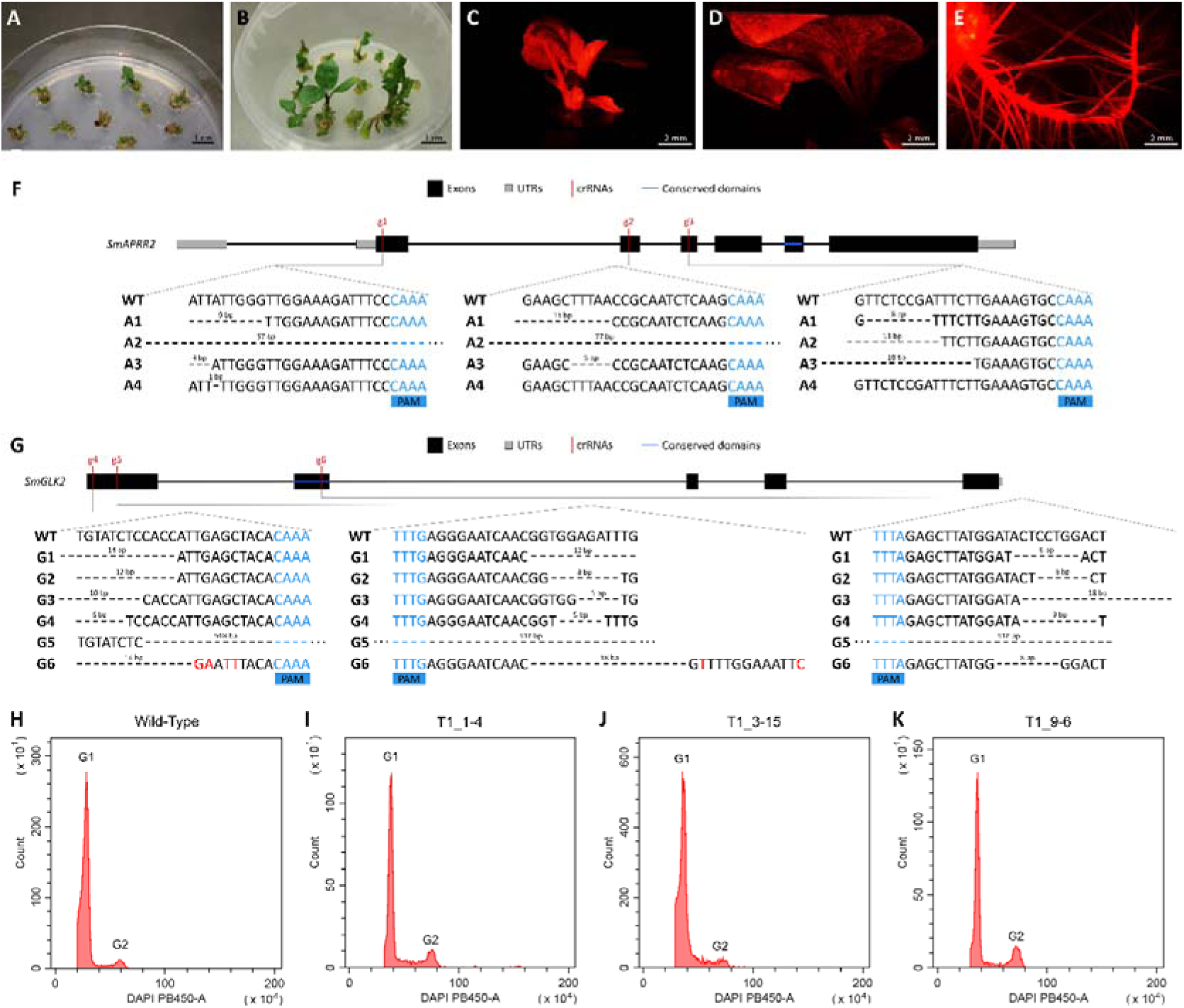
Recovery of transformed eggplant events, characterization of CRISPR/Cas12a-induced alleles, and flow-cytometric assessment of representative edited lines. (A, B) Adventitious shoot regeneration from cotyledon explants following *Agrobacterium*-mediated transformation, showing early regenerating shoots under selection (A) and elongated shoots with developed leaves (B). (C–E) Representative detection of *DsRed* fluorescence in a regenerated shoot (C), leaf (D), and roots (E). (F, G) Sequence characterization of the edited alleles identified in *SmAPRR2* (F) and *SmGLK2* (G). Gene models indicate exons (black), untranslated regions (UTRs; grey), and CRISPR RNA positions (crRNA, g1-g6, red). Wild-type (WT) target sequences and edited alleles are shown below for *SmAPRR2* (A1–A4) and *SmGLK2* (G1–G6). PAM sequences are highlighted in blue, deletions are represented by dashed gaps with their sizes indicated in base pairs, and nucleotide substitutions are shown in red. (H–K) Flow-cytometry histograms of nuclear DNA content in the WT (H) and representative edited T1 lines T1_1-4 (I), T1_3-15 (J), and T1_9-6 (K). The x-axis represents the fluorescence intensity emitted by DAPI and detected in the PB450 channel, expressed as the pulse area, which is proportional to nuclear DNA content. The y-axis indicates the number of detected nuclei for each fluorescence intensity level. The first peak corresponds to diploid nuclei in the G1 phase of the cell cycle. The second peak corresponds to diploid nuclei in the G2 phase, after DNA replication.

To assess segregation in the progeny, T_1_ seeds lacking *DsRed* fluorescence were selected, germinated, and genotyped by Sanger sequencing. PCR amplification of a *Cas12a* fragment confirmed that all selected T_1_ plants were transgene-free. Analysis of the transgene-free T_1_ progenies derived from the three founder events allowed the identification of homozygous, biallelic, and *APRR2^cr^/GLK2^cr^*plants (Table S2). Hereafter, *SmAPRR2*^cr^ and *SmGLK2*^cr^ denote genotypes carrying CRISPR-induced edited alleles at *SmAPRR2* and *SmGLK2*, respectively. Since T_0__1 carried edits exclusively in *SmGLK2*, 11 derived T_1_ individuals were analyzed (Table S2). Four of them (T_1__1-4, T_1__1-7, T_1__1-8, and T_1__1-10) were identified as *GLK2^cr^* lines, carrying two alternative mutation patterns in the three CRISPR RNAs (crRNAs) of *SmGLK2*, the −14/−12/−8 bp deletions (G1 allele) and −12/−8/−6 bp deletions (G2 allele), indicating that T_0__1 was biallelic (Figure 1G).

For the T_0__3-derived progeny, 29 T_1_ plants were genotyped (Table S2). Two distinct edited alleles were identified in *SmAPRR2,* namely the −9/−11/−8 bp deletions (A1 allele) and the −57/−77/−11 bp deletions (A2 allele), together with two edited alleles in *SmGLK2,* corresponding to the −10/−5/−18 bp deletions (G3 allele) and the −6/−5/−9 bp deletions (G4 allele) (Figure 1F, G). Based on these genotypes, five *APRR2^cr^/GLK2^cr^* plants (T_1__3-1, T_1__3-8, T_1__3-13, T_1__3-15, and T_1__3-20) were selected. In addition, four plants heterozygous for *SmGLK2* but homozygous for *SmAPRR2* (T_1__3-5, T_1__3-14, T_1__3-19, and T_1__3-23) were retained for further segregation analysis to recover *APRR2^cr^* lines.

T_1_ progeny derived from T_0__9, which was initially presumed to be homozygous for a large *SmGLK2* deletion of 918 bp (G5 allele), was also analyzed (Table S2). However, this subsequent analysis revealed that these plants were in fact biallelic, with the large deletion masking a second edited allele, corresponding to the G6 allele (−14/−18/−8 bp deletions together with six additional nucleotide substitutions, T>G, G>A, G>T, C>T, A>T, and T>C) (Figure 1G). In addition, in the T_1_ progeny, two edited alleles were identified in *SmAPRR2*, the A3 allele, corresponding to −3/−5/−18 bp deletions, and the A4 allele, corresponding to a −1 bp deletion at the crRNA1 target (Figure 1F). Four T_1_ plants from this family were selected (T_1__9-1, T_1__9-2, T_1__9-5, and T_1__9-6) for further studies. Although additional lines carrying the desired genotypes were identified, they were excluded from subsequent analyses because of poor seed germination.

### 2.2. Segregation-based recovery of desired mutant genotypes

To recover *APRR2^cr^* lines carrying wild-type (WT) alleles at *SmGLK2*, two complementary strategies were followed. First, T_2_ progenies derived from T_1__3-5 and T_1__3-14, initially expected to segregate for a WT *SmGLK2* allele, were analyzed. However, no WT homozygous individuals at *SmGLK2* were recovered in these families, and the segregation patterns indicated that both parental T_1_ lines carried two edited *SmGLK2* alleles rather than a WT and an edited allele (Table S3). Consistently, pseudo-F_1_ (pF1) progenies obtained by crossing T_1__3-5 and T_1__3-14 lines with WT plants were heterozygous at *SmGLK2*, confirming that the parental lines were biallelically edited at this locus (Table S3). Therefore, this strategy did not allow recovery of the desired *APRR2^cr^* genotype in a WT *SmGLK2* background.

As these initial approaches did not yield an *APRR2^cr^*line, an alternative strategy was implemented. Two *APRR2^cr^/GLK2^cr^*T_1_ lines (T_1__3-1 and T_1__3-15) and one *SmGLK2* biallelic line (T_1__9-5) were crossed with the WT. All resulting pF_1_ plants were heterozygous (Table S3) and were self-pollinated to generate pF_2_ populations (Figure S1). For each pF_1_-derived family, 90 pseudo-F_2_ (pF_2_) individuals were genotyped by Sanger sequencing (Table S4), and *APRR2^cr^* plants were selected for self-pollination. Assuming independent segregation, the expected frequency of this genotype combination was 1/16. Based on genotyping results, 12 plants with the desired genotype (*APRR2^cr^*) were selected for self-pollination. In addition, three extra plants were also selected as *APRR2^cr^/GLK2^cr^*lines, as they carried the alternative edited allele G6 for *SmGLK2* (Figure S1).

### 2.3. Characterization of CRISPR/Cas12a-induced indels in *SmGLK2* and *SmAPRR2*

Sequence analysis of edited plants revealed that CRISPR/Cas12a-induced mutations in both *SmAPRR2* and *SmGLK2* consisted predominantly of deletions affecting the regions targeted by the corresponding crRNAs (Figure 1F, G, Figure S2). In *SmGLK2*, edited alleles were detected in all 15 genotyped T_0_ plants, indicating a higher editing frequency at this locus than at *SmAPRR2*, where mutations were identified in nine plants (Table S1). Across the analyzed progenies, a limited number of recurrent edited alleles, corresponding to four *SmAPRR2* alleles and six *SmGLK2* alleles, was recovered, suggesting that several regenerants may have originated from common edited cell lineages.

For *SmAPRR2*, the predominant edited alleles corresponded to deletions of −9/−11/−8 bp (A1), −57/−77/−11 bp (A2), and -3/-5/-18 bp (A3), whereas an additional single-base deletion (−1 bp) allele (A4) was identified in the fixed progeny recovered after segregation (Figure 1F, Figure S2, Table S2). For *SmGLK2*, the most frequently recovered alleles consisted of small deletions affecting the three target sites, including the −14/−12/−8 bp (G1), −12/−8/−6 bp (G2), −10/−5/−18 bp (G3), −6/−5/−9 bp (G4), and −14/−18/−8 bp (with six extra nucleotide substitutions) (G6) profiles (Figure 1G, Figure S2). In addition, one edited event initially appeared to carry a large -918 bp deletion (G5), although segregation analysis later demonstrated that this line was in fact biallelic, with the large deletion masking the G6 edited allele (Figure 1G, Figure S2).

Overall, these results indicate that CRISPR/Cas12a editing generated stable and heritable deletion alleles in both target genes, ranging from small indels affecting individual target regions to larger deletions spanning broader genomic intervals. The recovery of different edited profiles provides the allelic basis for subsequent genotype–phenotype analyses.

### 2.4. Predicted effects on the protein sequence of representative edited alleles

To further assess the predicted functional impact of the edited alleles, the WT and mutant coding sequences of *SmAPRR2* and *SmGLK2* were translated *in silico* from the canonical start codon to the first in-frame stop codon. The predicted WT proteins were 378 aa and 234 aa in length for *SmAPRR2* and *SmGLK2,* respectively (Table 1). All edited alleles introduced premature termination events and were therefore predicted to encode truncated protein products relative to the corresponding WT sequences.

**Table 1.** Predicted effects of representative CRISPR/Cas12a-induced alleles on the SmAPRR2 and SmGLK2 proteins, including protein truncation, conserved domain retention, and putative functional consequences. WT and mutant coding sequences were translated *in silico* from the canonical start codon to the first in-frame stop codon to estimate the length of the predicted protein products. The percentage of WT protein retained was calculated relative to the corresponding WT sequence. Predicted consequences were inferred from the extent of protein truncation.

| Gene | Allele | Indel profile | WT protein length (aa) | Predicted mutant protein length (aa) | WT protein retained (%) | Conserved domain retained | Predicted consequence |
| --- | --- | --- | --- | --- | --- | --- | --- |
| SmAPRR2 | A1 | −9/−11/−8 | 378 | 65 | 17.2 | No | Severe truncation; likely loss-of-function. |
| SmAPRR2 | A2 | −57/−77/−11 | 378 | 298 | 78.8 | REC conserved domain putatively retained | N-terminally altered truncated protein; Putatively hypomorphic allele. |
| SmAPRR2 | A3 | −3/−5/−18 | 378 | 66 | 17.5 | No | Severe truncation; likely loss-of-function. |
| SmAPRR2 | A4 | −1 | 378 | 32 | 8.5 | No | Very severe truncation; likely strong loss-of-function. |
| SmGLK2 | G1 | −14/−12/−8 | 234 | 8 | 3.4 | No | Very severe truncation; likely strong loss-of-function. |
| SmGLK2 | G2 | −12/−8/−6 | 234 | 48 | 20.5 | No | Severe truncation; likely loss-of-function. |
| *SmGLK2* | G3 | -10/-5/-18 | 234 | 191 | 81.6 | *GLK2*<br>conserved<br>domain<br>putatively<br>retained | N-terminally altered<br>truncated protein;<br>Putatively hypomorphic<br>allele. |
| *SmGLK2* | G4 | -6/-5/-9 | 234 | 51 | 21.8 | No | Severe truncation; likely<br>loss-of-function. |
| *SmGLK2* | G5 | -918 | 234 | 7 | 3.0 | No | Very severe truncation;<br>likely strong loss-of-function. |
| *SmGLK2* | G6 | -14/-18/-8<br>T>G, G>A,<br>G>T, C>T,<br>A>T, T>C | 234 | 8 | 3.4 | No | Very severe truncation;<br>likely strong loss-of-function. |

In *SmAPRR2*, the severity of truncation varied markedly among alleles (Table 1). The shortest predicted product corresponded to A4 (32 aa), followed by A1 and A3, which encoded peptides of 65 aa and 66 aa, respectively. These three alleles were not predicted to retain the conserved domain and are therefore expected to have a strong disruptive effect on the encoded protein. The allele A2 retained a larger portion of the protein, with a predicted length of 298 aa. In this case, the conserved domain was putatively retained. Alignment with the WT protein showed that the main differences were confined to the N-terminal region, whereas the remaining sequence was highly similar to WT, suggesting that, if the transcript is not degraded by nonsense-mediated decay, A2 may therefore represent a putatively hypomorphic allele with partial residual functionality.

A similar pattern was observed in *SmGLK2*, where the alleles G1, G5, and G6 were predicted to encode extremely short peptides of 8 aa, 7 aa, and 8 aa, respectively, whereas G2 and G4 encoded truncated proteins of 48 aa and 51 aa (Table 1). None of these alleles was predicted to retain the conserved domain. By contrast, G3 retained a substantially longer predicted product of 191 aa, with the conserved domain putatively retained. Sequence alignment with the WT revealed that, aside from the N-terminal region affected by the mutation, the remaining predicted amino acid sequence was highly similar to that of the WT protein. Therefore, provided that the transcript is not degraded by nonsense-mediated decay, G3 could also represent a putatively hypomorphic allele with partial residual functionality.

Overall, these results indicate that most analyzed alleles are expected to have a strong disruptive effect on the encoded proteins. However, the longer predicted products encoded by A2 and G3 may retain some degree of biological activity, as their conserved domains were putatively retained and their predicted sequences remained highly similar to the WT outside the mutated N-terminal region.

### 2.5. Phenotypic characterization of selected fixed edited lines

Selected homozygous edited lines carrying distinct allelic combinations at *SmAPRR2* and *SmGLK2* were evaluated for fruit color phenotypes (Table 2; Figure 2; Figure S3). These lines covered the three expected phenotypic classes derived from the single- and double-gene mutant combinations: green fruits without visible netting (from *GLK2^cr^* lines), fruits with netting (from *APRR2^cr^* lines), and fully white fruits (from *APRR2^cr^/GLK2^cr^*lines). At the whole-plant level, no obvious differences in vegetative development or leaf green pigmentation were visually observed between the edited lines and WT plants under the evaluated greenhouse conditions.

**Figure 2.**
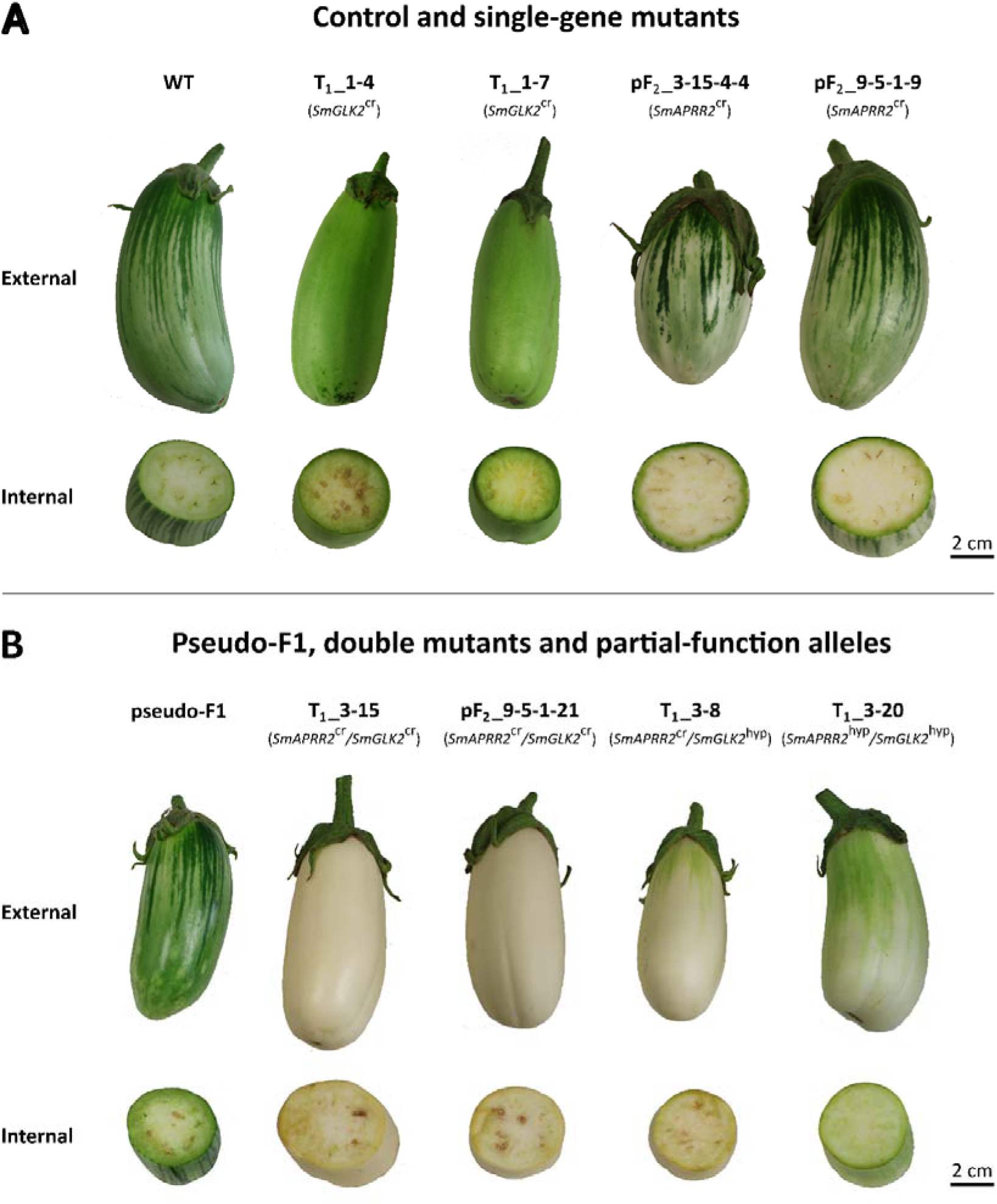
External and internal fruit pigmentation phenotypes associated with different *SmAPRR2* and *SmGLK2* allele combinations. External fruit views are shown in the upper row of each panel, with the corresponding transverse sections directly below. (A) Wild type (WT) and single-gene edited lines: *SmGLK2*^cr^ lines T_1__1-4 and T_1__1-7, and *SmAPRR2*^cr^ lines pF_2__3-15-4-4 and pF_2__9-5-1-9. (B) Pseudo-F_1_, double-mutant lines (*SmAPRR2*^cr^*/SmGLK2*^cr^) T_1__3-15 and pF_2__9-5-1-21, and lines carrying predicted hypomorphic alleles T_1__3-8 and T_1__3-20. Scale bars = 2 cm. cr: CRISPR-induced loss-of-function allele. hyp: predicted hypomorphic allele retaining partial function.

**Table 2.**
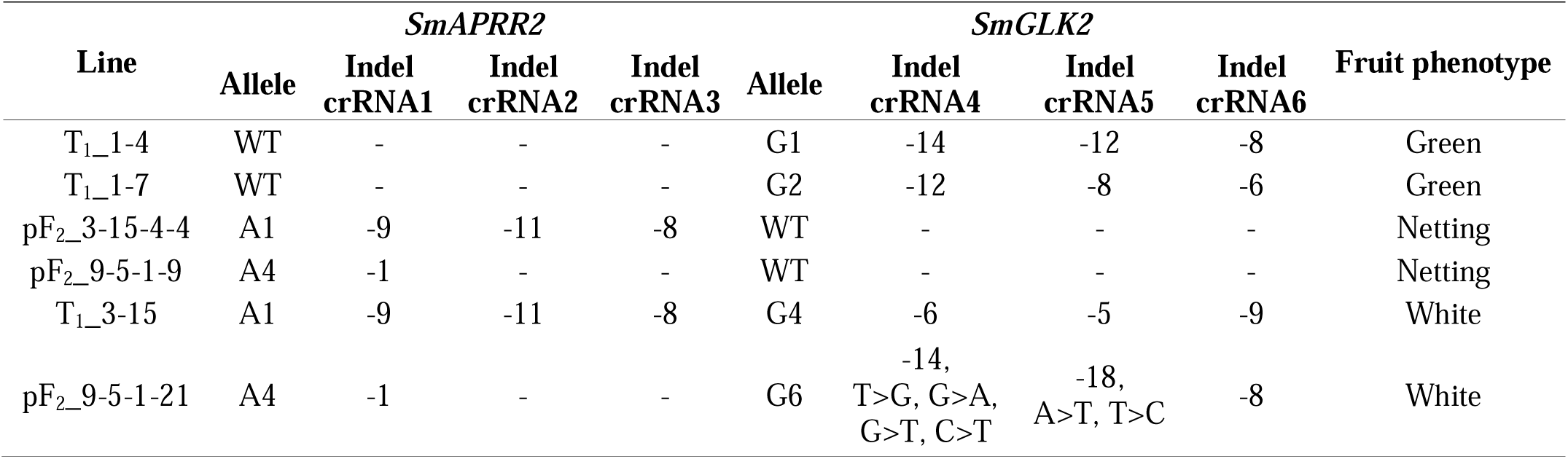
Fixed edited lines selected for phenotypic analysis, representing the three fruit color phenotypes and their corresponding *SmAPRR2* and *SmGLK2* indel profiles for each CRISPR RNA (crRNA1-6).

Fruits from WT and pF_1_ plants showed visible green pigmentation externally (Figure 2). *GLK2^cr^* lines with putative loss of function (e.g., T_1__1-4 and T_1__1-7) produced fruits with a uniform green phenotype, in which the green netting pattern was absent (Figure 2A). This supports the role of *SmGLK2* in the formation of the reticulated chlorophyll pigmentation pattern in the fruit peel. In contrast, *APRR2^cr^*lines with putative loss of function (e.g., pF_2__3-15-4-4 and pF_2__9-5-1-9) displayed fruits with dark green netting over a light green background (Figure 2A), indicating that *SmAPRR2* contributes mainly to the overall level of chlorophyll pigmentation in the fruit, while *SmGLK2* is primarily involved in green netting formation but also appears to contribute to the green background coloration. *APRR2^cr^/GLK2^cr^* lines carrying disruptive mutations in both *SmAPRR2* and *SmGLK2* (e.g., T_1__3-15 and pF_2__9-5-1-21) produced fully white fruits with no visible green pigmentation (Figure 2B), consistent with the combined disruption of both chlorophyll-related pathways.

Interestingly, two *APRR2^cr^/GLK2^cr^* lines, T_1__3-8 and T_1__3-20, showed milder pigmentation phenotypes than the fully white double mutants (Figure 2B). T_1__3-8 carried the *SmGLK2* G3 and the *SmAPRR2* A1 alleles and produced nearly white fruits, but residual green pigmentation remained mainly restricted to the proximal region and to small areas of the fruit surface. This phenotype is consistent with the predicted partial residual functionality of the G3 allele (Table 1). Similarly, T_1__3-20 carried the *SmGLK2* G3 and the *SmAPRR2* A2 alleles and it showed a less severe phenotype, retaining more evident diffuse green pigmentation over the fruit surface, in agreement with the predicted residual functionality of G3 and A2 allele variants (Table 1). These observations suggest that not all edited alleles at *SmGLK2* and *SmAPRR2* necessarily behave as complete loss-of-function mutations, and that some of them could represent putatively hypomorphic alleles associated with residual gene activity that can partially maintain chlorophyll-related pigmentation.

Cross-sections of commercially immature fruits further showed that the external pigmentation phenotypes were accompanied by consistent changes in internal chlorophyll-related pigmentation (Figure 2). WT, pF1, and *SmGLK2*^cr^ fruits displayed a general green peel pigmentation, a clearly visible green ring in the flesh immediately beneath the fruit peel together with a green peripheral region and a white central parenchyma (Figure 2). This indicates that disruption of *SmGLK2*, although sufficient to alter the external green netting pattern, did not abolish the internal green ring.

By contrast, *APRR2^cr^* fruits lost the internal green ring, although the tissue closest to the peel remained green, indicating that chlorophyll pigmentation was retained only in the outermost fruit flesh, due to the presence of the green netting pattern (Figure 2A). Finally, *APRR2^cr^/GLK2^cr^*fruits lacked visible green pigmentation in the inner tissues, showing a complete loss of the green ring (Figure 2B). Among the double-edited lines with milder external pigmentation phenotypes, T_1__3-8 also showed a white internal phenotype in the section analyzed. By contrast, T_1__3-20 retained internal green pigmentation, including a visible green ring and green peripheral tissue. These observations support a major role for *SmAPRR2* in maintaining chlorophyll accumulation in both the fruit peel and the flesh-adjacent green ring, whereas *SmGLK2* appears to contribute mainly to the green pigmentation of the outermost flesh adjacent to the peel, likely associated with the external netting pattern.

### 2.6. Off-target analysis

Whole-genome resequencing data showed high overall alignment quality for all analyzed samples. Total reads ranged from 265,135,844 to 293,209,832, the percentage of mapped reads ranged from 97.82% to 98.59%, and the percentage of mapped reads ranged from 97.82% to 98.59% (Table S5). On-target coverage was also high for both target genes, with mean coverage values of 99.56% for *SmAPRR2* and 98.86% for *SmGLK2* across the evaluated sample-gene combinations (Table S6).

The TTTV ≤ 4-mismatch sensitivity search identified one on-target hit for each of the six crRNAs and only two predicted off-target sites across the genome (Table S7). One predicted off-target corresponded to *SmAPRR2* crRNA1 on chromosome 6 and contained two mismatches relative to the guide sequence. The second predicted off-target corresponded to *SmGLK2* crRNA6 on chromosome 9 and contained four mismatches. No predicted off-target sites were identified for *SmAPRR2* crRNA2, *SmAPRR2* crRNA3, *SmGLK2* crRNA4, or *SmGLK2* crRNA5.

Both predicted off-target sites were fully covered in all seven analyzed samples, including the WT (*S. melongena* accession MEL3) and the six edited lines (Table S8). The *SmAPRR2* crRNA1 candidate site showed a mean depth of 35.16× across samples, with a minimum depth of 26.34×, whereas the *SmGLK2* crRNA6 candidate site showed a mean depth of 34.36×, with a minimum depth of 27.99×. In both cases, mean, minimum, and maximum coverage across the protospacer ± 100 bp interval were 100%, and no zero-depth bases were detected.

Variant inspection did not identify any candidate off-target variant absent from MEL3 reference at either predicted site. In particular, no candidate indels were detected within the protospacer ± 100 bp intervals, and none overlapped the protospacer sequences themselves. Therefore, under the TTTV ≤4-mismatch sensitivity analysis, no evidence of Cas12a-induced off-target editing was detected in the selected edited lines.

### 2.7. Flow cytometry analysis

Ploidy analysis by flow cytometry showed that all analyzed T_1_ plants displayed a diploid DNA content profile, with G1 and G2 peaks at positions comparable to those of the WT. Representative histograms for the WT and one selected T_1_ line derived from each of the three independent T_0_ events are shown in Figure 1H-K, and the complete set of histograms for all analyzed T_1_ plants is provided in Figure S4. The absence of shifts in peak position relative to the WT indicated that no ploidy alterations were detected in any of the evaluated T_1_ lines. These results confirmed that the selected edited progenies retained the expected diploid ploidy level.

## 3. Discussion

This study establishes CRISPR/Cas12a-mediated gene editing in eggplant and uses this platform to dissect the complementary roles of *SmAPRR2* and *SmGLK2* in chlorophyll-related pigmentation of the fruit peel and flesh. To our knowledge, this is the first demonstration of Cas12a-mediated editing in eggplant, expanding the genome-editing toolkit available for this recalcitrant crop (Maioli et al., 2020; Wang et al., 2022; Kodackattumannil et al., 2023; Phad et al., 2024; Sagarbarria et al., 2023; Ferrero et al., 2024, 2026; Liao et al., 2024; Zhang, Zhang, Yan et al., 2024). Previous studies had identified *SmAPRR2* as a strong candidate for uniform chlorophyll pigmentation in the fruit peel and *SmGLK2* as the candidate underlying the irregular green netting phenotype through genetic mapping, association analyses, and expression data (Arrones et al., 2022, 2024). Here, targeted mutagenesis of these loci allowed us to move from candidate-gene association to functional demonstration. Importantly, this validation was supported by the recovery of several independent edited alleles and fixed genotype combinations, strengthening the causal link between each target gene and the corresponding fruit pigmentation phenotype. Thus, our work bridges genetic mapping, allele engineering, and phenotypic validation, while providing a proof-of-concept for CRISPR/Cas12a-mediated functional genomics and precision trait design in eggplant.

The edited allelic combinations allowed chlorophyll-related fruit coloration to be dissected into separable phenotypic components. Disruption of *SmAPRR2* reduced the overall green background of the fruit peel while retaining the netting pattern, indicating that *SmAPRR2* acts primarily as a major determinant of chlorophyll accumulation intensity. This finding places *SmAPRR2* as a central regulator of background chlorophyll pigmentation in eggplant fruit, rather than as a general modifier of the fruit color pattern (Arrones et al., 2022; Fang et al., 2023). This interpretation agrees with the conserved role of *APRR2-like* genes in plastid development and fruit pigmentation across horticultural crops. In tomato, overexpression of an *APRR2-like* gene increased chlorophyll content in unripe fruits, as well as plastid number and plastid area, while in pepper, variation in an orthologous *APRR2-like* gene was associated with differences in immature fruit pigmentation (Pan et al., 2013). Similarly, in cucurbits, *APRR2* homologues have been identified as causal or strong candidate genes for rind or peel green pigmentation in cucumber, melon, watermelon, pumpkin, and chayote, further supporting the recurrent recruitment of this regulatory module in horticultural fruit color diversification (Jiao et al., 2017; Oren et al., 2019; Gebretsadik et al., 2024; Cheng et al., 2025). More recently, functional analyses in cucumber showed that *CsAPRR2* coordinates chloroplast biogenesis and photosynthesis-related gene expression, with reduced *CsAPRR2* activity leading to impaired thylakoid development, lower chloroplast number, and reduced pigment content (Liu et al., 2025). Thus, our functional validation of *SmAPRR2* in eggplant extends a conserved *APRR2*-related pigmentation module to Solanaceae fruit color diversification and identifies this gene as a promising target for precision modulation of chlorophyll-related fruit pigmentation.

Disruption of *SmGLK2* eliminated the green netting pattern while maintaining the green background, indicating that *SmGLK2* primarily contributes to the spatial patterning of chlorophyll rather than to the peel’s overall capacity to uniformly accumulate green pigment. This places *SmGLK2* in a different functional layer from *SmAPRR2:* whereas *SmAPRR2* controls the intensity or presence of green pigmentation, *SmGLK2* determines how this pigmentation is distributed across the fruit surface. This interpretation is consistent with the conserved role of GLK transcription factors in chloroplast development and fruit chlorophyll patterning (Arrones et al., 2024). In tomato, *SlGLK2* underlies the green shoulder phenotype and variation at the Uniform Ripening locus affects chloroplast development, fruit appearance, and ripening-related quality traits (Powell et al., 2012; Nguyen et al., 2014). Additional evidence from tomato indicates that *SlGLK2* is embedded within a broader regulatory network controlling fruit chloroplast development, as the transcription factor *SlBEL2* negatively regulates green shoulder formation by repressing *SlGLK2* expression and interfering with *SlGLK2* protein activity (Niu et al., 2022). In pepper, *CaGLK2* underlies natural variation in chlorophyll content and immature fruit color, and recent evidence shows that *GLK2* and *PRR2/APRR2*-related regulators can jointly modulate fruit pigmentation within a shared plastid-pigmentation module (Brand et al., 2014; Lee et al., 2026). Together, these studies indicate that GLK2-related genes contribute to chlorophyll-dependent pigmentation in immature fruits across Solanaceae (Brand et al., 2014; Nguyen et al., 2014; Arrones et al., 2024), but with different spatial manifestations depending on the crop. In tomato and eggplant, *GLK2* is associated with localized green pigmentation in the upper or proximal region of the fruit, appearing as the green shoulder in tomato (Powell et al., 2012) and as green netting in eggplant (Arrones et al., 2024). By contrast, in pepper, *CaGLK2* is more broadly associated with chlorophyll content and the overall green coloration of immature fruits (Lee et al., 2026).

The *APRR2^cr^/GLK2^cr^* lines were particularly informative because they revealed how both regulators interact to generate the final fruit pigmentation phenotype. In a WT *SmAPRR2* background, disruption of *SmGLK2* removed the netting pattern but did not eliminate the green background, indicating that *SmAPRR2* is sufficient to maintain strong uniform peel chlorophyll accumulation. Conversely, when *SmAPRR2* was disrupted, the contribution of *SmGLK2* became visible mainly as residual netting over a much lighter background. For background green coloration, these results suggest that *SmAPRR2* is epistatic to *SmGLK2*, as the contribution of *SmGLK2* to basal green coloration becomes mainly evident when *SmAPRR2* is disrupted (Dwivedi et al., 2024). Recent evidence in pepper provides an interesting parallel, as combined variation in *GLK2* and *PRR2/APRR2*-related regulators has been associated with pale-green immature fruits, supporting the idea that these genes can act within a shared plastid-pigmentation module in Solanaceae fruits (Lee et al., 2026). In our study, only the combined disruption of both genes resulted in fully white fruits, demonstrating that the most extreme loss of green pigmentation requires removal of both the green uniform background and netting components.

This genetic architecture has important implications for breeding by design (Zsögön et al., 2018). Rather than treating eggplant fruit color as a single qualitative trait, our results show that it can be decomposed into editable modules that can be recombined to generate predictable phenotypic outputs: green fruits without netting, light green fruits with netting, white fruits, and intermediate pigmentation phenotypes. This opens the possibility of manipulating the eggplant fruit color palette through targeted allele combinations, offering a route toward precision breeding and predictable valuation of visual quality traits (Arrones et al., 2022, 2024).

The analysis of fruit flesh green pigmentation further extends the functional role of *SmAPRR2* beyond the fruit surface. A major QTL for the presence of a green ring in the flesh next to the skin was previously mapped at the top of chromosome 8, where a ferredoxin-family gene was initially proposed as a candidate gene for this trait because of its potential involvement in chlorophyll production (Portis et al., 2014). However, the associated marker lies only 623 kb from *SmAPRR2*, which was later identified as a major candidate for fruit peel chlorophyll pigmentation (Arrones et al., 2022). Our edited materials now provide functional support for this genomic association: disruption of *SmAPRR2* strongly reduced the green ring, whereas *GLK2*^cr^ lines retained this internal pigmentation. This is consistent with evidence from cucurbit fruits showing that *APRR2*-related regulation can influence pigment accumulation and chloroplast development in internal fruit tissues, including the flesh (Oren et al., 2019; Liu et al., 2025; Valverde et al., 2025). Therefore, the green ring is more closely linked to *SmAPRR2*-dependent chlorophyll accumulation than to the *SmGLK2*-dependent external netting pattern. This refines the genetic model of eggplant fruit coloration (Arrones et al., 2022, 2024; Yang et al., 2022; You et al., 2023) by connecting peel pigmentation and flesh-adjacent chlorophyll accumulation through a shared regulatory component.

An additional strength of this work is the recovery of an informative edited allelic series. Most alleles were predicted to produce severe truncations and were associated with strong pigmentation phenotypes, consistent with major loss of function. However, T1_3-20 and T1_3-8 showed intermediate phenotypes, which were consistent with the presence of longer predicted protein products for the *SmAPRR2* A2 or *SmGLK2* G3 alleles, respectively. These putatively hypomorphic alleles indicate that CRISPR/Cas12a editing not only generated complete loss-of-function mutations, but also alleles with partial residual activity. From a breeding perspective, these intermediate phenotypes are particularly valuable because they illustrate that genome editing can, aside from generating complete knockouts, create *de novo* allelic variation for fine-tuning fruit quality traits, enabling gradations of pigmentation beyond a simple green-versus-white contrast (Yoshimura and Ishida, 2025). Such allele engineering could be especially useful for trait tailoring, where subtle modulation of color intensity, pattern visibility, or internal pigmentation may be preferable to complete loss of pigment (Wolter et al., 2019).

Beyond pigment biology, our results expand the genome-editing toolkit available for eggplant by introducing Cas12a as an efficient editing platform in this crop. Previous genome-editing studies in this species have relied on CRISPR/Cas9 and have mainly focused on reducing enzymatic browning through *PPO* gene editing, establishing or evaluating editing systems using targets such as *SmWRKY4* and *SmPDS*, and validating genes involved in agronomic or developmental traits, including anthocyanin accumulation, *Phytophthora* tolerance, parthenocarpy, and prickle formation (Maioli et al., 2020; Wang et al., 2022; Kodackattumannil et al., 2023; Phad et al., 2024; Sagarbarria et al., 2023; Ferrero et al., 2024; Liao et al., 2024; Zhang, Zhang, Yan et al., 2024; Toppino et al., 2025). In addition, CRISPR/Cas9 has recently been applied to species other than *S. melongena* within the eggplant genepool, including wild and cultivated *Solanum* relatives, to functionally validate LOG cytokinin biosynthetic genes involved in prickle development and to generate prickleless edited plants (Satterlee et al., 2024). Therefore, our study provides a proof-of-concept for Cas12a-mediated editing in eggplant and demonstrates that this nuclease can generate stable, heritable alleles, and transgene-free progenies after segregation in eggplant.

Cas12a offers features particularly attractive for plant genome editing, including a T-rich PAM, staggered DNA cleavage, and straightforward multiplexing capability (Banakar et al., 2020; Bandyopadhyay et al., 2020; Paul and Montoya, 2020). The value of Cas12a as an alternative to Cas9 has already been demonstrated in tomato, where LbCas12a was shown to be an efficient and robust nuclease, with target-dependent activities comparable to SpCas9 and a tendency to generate larger deletions, a feature that may be advantageous when complete gene disruption is desired (Slaman et al., 2024). Cas12a-derived platforms have also been applied in tomato for precise genome editing and transcriptional activation, including LbCpf1/Cas12a-based gene targeting and dCas12a-mediated upregulation of *SlPAL2* to enhance resistance to bacterial canker (Vu et al., 2020, 2021; Rivera-Toro et al., 2025). In this context, the successful implementation of Cas12a in eggplant represents a meaningful expansion of the editing toolbox for a crop in which functional genomics has historically been constrained by transformation and regeneration bottlenecks.

The resequencing-based off-target assessment further supports the suitability of Cas12a for functional validation in eggplant. Under a conservative TTTV ≤ 4-mismatch search, only two predicted off-target sites were identified for the six crRNAs, and both were completely covered in the resequencing data. The absence of candidate variants or indels absent from MEL3 at these sites indicates that the selected edited lines did not show detectable sequence changes compatible with Cas12a-induced off-target editing at the predicted loci. This result is particularly relevant because specificity is a key consideration when introducing Cas12a as an alternative editing platform in a crop where functional genomics remains technically challenging (Bandyopadhyay et al., 2020). This is consistent with previous reports showing that Cas12a generally displays stringent target recognition and relatively low mismatch tolerance, particularly in PAM-proximal regions, which contributes to its high editing specificity (Kim et al., 2020; Murugan et al., 2020). At the same time, the analysis should be interpreted within its defined search space: it provides strong evidence against editing at predicted TTTV-associated off-target sites with up to four mismatches, but it does not replace broader whole-genome structural variant analyses or assays targeting non-canonical PAM contexts in larger edited panels (Graham et al., 2020; Sturme et al., 2022). In this context, whole-genome resequencing has been used as a powerful approach to assess potential off-target mutations in plants (Tang et al., 2018; Bernabé-Orts et al., 2019; Slaman et al., 2024).

Nevertheless, the low number of independent edited events confirms that transformation and regeneration remain major bottlenecks in this species. Eggplant is more recalcitrant than tomato (Saini et al., 2025), and previous reports have emphasized the strong influence of genotype, explant type, regeneration capacity, and transformation conditions on editing success (García-Fortea et al., 2020; Khatun et al., 2022; Sagarbarria et al., 2023). In our case, extensive segregation and crossing schemes were required to recover the desired allelic combinations, illustrating the practical complexity of functional validation in eggplant. However, all genotyped transgenic regenerants carried CRISPR-induced mutations, suggesting that once transformation was achieved, the Cas12a editing system was highly effective and efficient in the recovered events. Thus, the main limitation of the workflow for eggplant gene editing appears to be the recovery of independent transformed regenerants rather than editing activity itself. Future improvements in regeneration protocols, genotype-independent transformation, and multiplex editing design will be essential for moving towards standardized genome-editing workflows and making editing in recalcitrant crops such as eggplant more routine (Cardi et al., 2023).

Overall, this work establishes CRISPR/Cas12a as a functional genome-editing platform for eggplant and demonstrates its utility for candidate-gene validation, predictable trait modulation, and the generation of putatively hypomorphic alleles that create new phenotypic variation. Together with the absence of detectable editing at the two predicted TTTV-associated off-target sites, these findings support the use of Cas12a for targeted functional genomics in eggplant, while remaining restricted to the off-target search space and edited panel evaluated here. By separating chlorophyll-related pigmentation into background coloration, green netting, and internal green-ring components, the study provides a refined genetic model for eggplant fruit color and shows how targeted allele combinations can be used to engineer distinct visual phenotypes. More broadly, our results illustrate how genome editing can convert genetic associations into validated breeding targets, expand the editable phenotypic space of recalcitrant crops, and enable precision manipulation of horticultural quality traits. In this sense, our work contributes to the development of molecular horticulture in eggplant by linking genetic mapping, Cas12a-mediated allele engineering, and visible quality traits in a crop where functional validation remains comparatively underdeveloped.

## 4. Materials and methods

### 4.1. Plant material

The MEL3 accession of *S. melongena*, provided by the germplasm bank at Universitat Politècnica de València (Valencia, Spain; FAO germplasm bank code: ESP026), was used in this study. The MEL3 accession exhibits fruits with a uniform green peel background combined with an irregular dark green netting pattern (García-Fortea et al., 2020) and was characterized both morphologically and genetically (Kaushik et al., 2016, 2017), confirming that it carries functional alleles of both *SmAPRR2* and *SmGLK2*.

### 4.2. Candidate genes

The *SmAPRR2* gene (SMEL_008g315370) and *SmGLK2* gene (SMEL_004g203570), as annotated in the eggplant reference genome “67/3” v3.0 (Barchi et al., 2019), were selected for functional analysis. *SmAPRR2* is located on chromosome 8, while *SmGLK2* is located on chromosome 4 (Figure 3A). *SmAPRR2* contains a conserved phosphoacceptor receiver (REC) domain of response regulators (RRs) and pseudo response regulators (PRRs) in exon 5, whereas exon 2 of *SmGLK2* contains the conserved golden-2 like transcription factor, PLN03162 Superfamily domain (Figure 3A). The 5′ genomic regions encompassing the selected target exons of *SmAPRR2* and *SmGLK2* were amplified from MEL3 genomic DNA by PCR using the primer pairs APRR2_F/APRR2_R, and GLK2_F/GLK2_R, respectively (Table S9). These gene-specific primers were designed using Primer3 (https://primer3.ut.ee/). Specifically, for *SmAPRR2*, exons 1, 2, and 3 were amplified, whereas for *SmGLK2*, exons 1 and 2 were targeted. The amplified fragments were then subjected to Sanger sequencing to identify MEL3-specific sequence variants relative to the reference genome from SolGenomics, ensuring the accuracy of the sequence for subsequent crRNA design.

**Figure 3.**
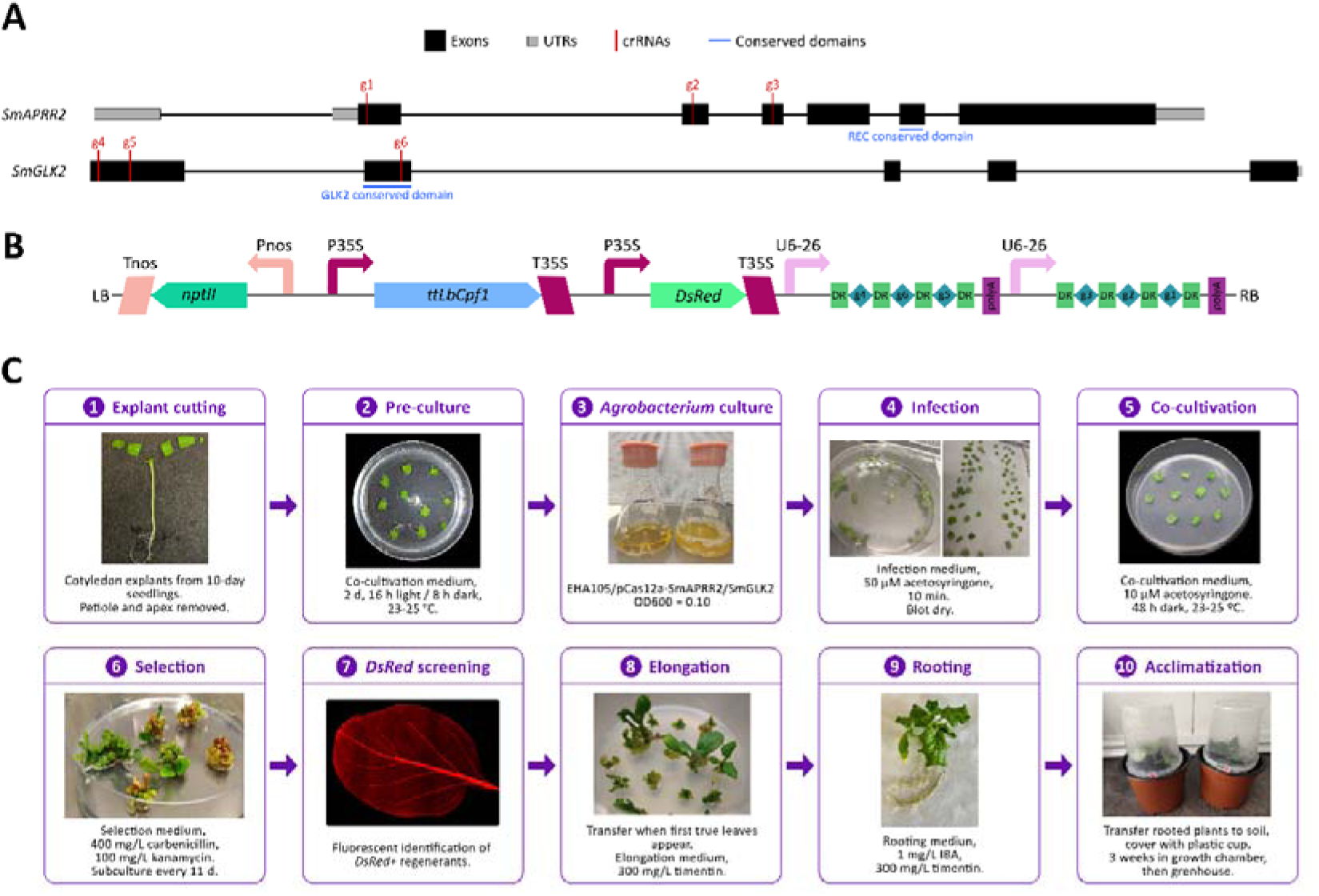
CRISPR/Cas12a editing strategy and *Agrobacterium*-mediated transformation workflow for *SmAPRR2* and *SmGLK2* genes in eggplant. (A) Gene structures of *SmAPRR2* and *SmGLK2*, showing exons in black, untranslated regions (UTRs) in grey, and conserved domains indicated by blue lines. The positions of the designed crRNAs are indicated by red vertical lines. (B) Genetic construct assembled using the GoldenBraid system and used for CRISPR/Cas12a-mediated editing of *SmAPRR2* and *SmGLK2* genes. (C) Overview of the *Agrobacterium*-mediated transformation and regeneration workflow in eggplant. The figure summarizes the main steps of the protocol, from cotyledon explant preparation to acclimatization of transformed plants. Culture media used at each stage correspond to those described in Table S11. LB: left border. Tnos: nopaline synthase terminator. *nptII*: *neomycin phosphotransferase II*. Pnos: nopaline synthase promoter. P35S: cauliflower mosaic virus 35S promoter. ttLbCpf1: temperature-tolerant *Lachnospiraceae bacterium* Cpf1/Cas12a. T35S: cauliflower mosaic virus 35S terminator. *DsRed*: *Discosoma* red fluorescent protein reporter gene. U6-26: *Arabidopsis thaliana* U6-26 promoter. DR: Cas12a direct repeat. g1, g2, g3, g4, g5 and g6: CRISPR RNAs (crRNAs) (Table S10). RB: right border.

### 4.3. Plasmid design and construction

Specific crRNAs targeting the *SmAPRR2* and *SmGLK2* genes were designed based on the corresponding Sanger sequences obtained from MEL3. The process involved a comprehensive analysis using several computational tools to ensure optimal targeting efficiency and specificity. CRISPOR (https://crispor.gi.ucsc.edu/), CCTop (https://cctop.cos.uni-heidelberg.de/), and Benchling (https://www.benchling.com/) tools were used to select crRNAs with high predicted on-target activity and minimal off-target potential. For *SmAPRR2*, three crRNAs were designed: crRNA1 in exon 1, crRNA2 in exon 2, and crRNA3 in exon 3 (Figure 3A, Table S10). Similarly, for *SmGLK2*, three crRNAs were designed: crRNA4 and crRNA5 targeting exon 1, and crRNA6 targeting exon 2, within the GLK2 conserved domain (Figure 3A, Table S10).

The crRNAs were synthesized and assembled into a CRISPR/Cas12a construct carrying the *neomycin phosphotransferase II* (*nptII)*, *Cas12a (LbCpf1),* and *DsRed* genes (Figure 3B), using the GoldenBraid (GB) assembly system (https://goldenbraidpro.com/), following the guidelines provided by GB software (Vazquez-Vilar et al., 2017). To ensure compatibility with the GB system, the crRNA sequences and their corresponding Cas12a direct repeats (DRs) were domesticated, and a DNA fragment was synthesized by GenScript company (https://www.genscript.com/). The CRISPR construct was designed to simultaneously target both genes, so the *SmAPRR2* crRNAs were arranged in a polycistronic crRNA array driven by the AtU6-26 RNApol III promoter, while the *SmGLK2* crRNAs were expressed using a separate promoter. The expression of *Cas12a* and *DsRed* genes was regulated by the CaMV 35S promoter, with the T35S terminator, while the *nptII* gene was expressed under the control of the Pnos promoter, coupled with the Tnos terminator (Figure 3B). The final vector, designated pCas12a-SmAPRR2/SmGLK2, was introduced into electrocompetent *Agrobacterium tumefaciens* strain EHA105 by electroporation. Briefly, 1 µL of plasmid DNA was mixed with 50 µL of electrocompetent EHA105 cells in a pre-chilled electroporation cuvette and pulsed at 1440 V. Immediately after electroporation, 500 µL of SOC medium was added, and cells were recovered for 2 h at 28 °C with shaking at 200 rpm. Transformants were selected on LB agar plates containing rifampicin and kanamycin and incubated at 28 °C for two days. The resulting recombinant strain was subsequently referred to as EHA105/pCas12a-SmAPRR2/SmGLK2.

### 4.4. *Agrobacterium*-mediated transformation

The eggplant transformation procedure followed previously described protocols (Rotino and Gleddie, 1990; Arpaia et al., 1997; Rotino et al., 1997; Toppino et al., 2011, 2025; Florio et al., 2021), with several modifications (Figure 3C). For *in vitro* germination, seeds of MEL3 were surface-sterilized by immersion in 70% ethanol for 30 s, followed by treatment with a 20% (v/v) commercial bleach solution (with 3.8% NaClO) for 15 min. Subsequently, the seeds were rinsed three times with sterile deionized water for 3–5 min each (García-Fortea et al., 2020). After sterilization, seeds were soaked in a 500 ppm GA□ solution for 24 h to promote germination (Ranil et al., 2015). The following day, the seeds were placed on germination medium (Table S11) and cultured in Microbox containers (9.2 cm diameter × 12 cm height) equipped with a membrane filter in the lid to allow gas exchange (O118/120+OD118/120 #10 [NG/NP], SAC02, Nevele, Belgium).

For transformation experiments (Figure 3C), the *Agrobacterium tumefaciens* strain EHA105 carrying the construct pCas12a-SmAPRR2/SmGLK2 (EHA105/pCas12a-SmAPRR2/SmGLK2) was prepared as follows. Two days prior to infection, bacteria were revived from a glycerol stock and grown for 28 h at 28 °C with agitation (180 rpm) in 5 mL of LB medium supplemented with kanamycin and rifampicin (50 mg/L each). This pre-culture was subsequently used to inoculate 100 mL of TY medium (5 g/L tryptone, 3 g/L yeast extract, 203.3 mg/L MgCl□·6H□O and 246.5 mg/L MgSO□·7H□O, pH 5.8) supplemented with 50 mg/L kanamycin, with different inoculum volumes (2.5, 5, 10, and 15 µL) to adjust the optical density at 600 nm (OD□□□) more easily. Cultures were incubated for 16 h at 28 °C with shaking at 200 rpm until an OD□□□ of approximately 0.10 was reached. Bacterial cells were then harvested by centrifugation at 4,000 × g for 5 min at room temperature, and the resulting pellet was resuspended by vortexing in an equal volume of infection medium (Table S11).

Ten days after sowing the seeds on germination medium, and prior to the emergence of the first true leaves, cotyledon explants were excised two days before infection by removing the petioles and apices (Figure 3C). When cotyledons were sufficiently large, they were divided into two explants. In all culture steps, explants were placed with the abaxial side down. The explants were placed on co-cultivation medium (Table S11) and incubated for two days in a growth chamber under a 16 h light/8 h dark photoperiod (100–112 µmol m□² s□¹) at 23–25 °C. Culture plates were sealed with Micropore tape throughout the *in vitro* culture process.

For infection, explants were immersed in the *Agrobacterium* inoculum prepared in infection medium for 10 min and subsequently blotted on sterile filter paper to remove excess bacteria (Figure 3C). The explants were then returned to the co-cultivation medium and maintained in darkness at 23–25 °C for 48 h. After co-cultivation, explants were transferred to the selection medium (Table S11) and subcultured onto freshly prepared medium every 14 days until callus formation. Proliferating calli were removed from explants, cleaned from dead tissue and subcultured onto freshly prepared medium until green shoots-buds begin to form. Once the regenerated shoots developed the first true leaves, they were transferred to the elongation medium (Table S11). When the shoots reached the plate lid, they were excised and transferred to rooting medium (Table S11) in Microbox containers, where they were cultured until root formation was observed. For acclimatization, well-rooted plantlets were washed to remove the culture medium and transferred to soil, where they were initially covered with plastic cups that were gradually removed (Figure 3C). Plants were maintained in a growth chamber for three weeks before transferring them to the greenhouse.

Transformation efficiency was calculated as the percentage of initially infected explants that produced transgenic regenerants, according to the formula: transformation efficiency (%) = (number of explants producing transgenic regenerants / total number of infected explants) × 100. The 95% confidence interval for the observed proportion was calculated using the exact binomial method.

### 4.5. Molecular characterization and selection of edited lines

Genomic DNA from T_0_ plants was isolated from young leaf tissue using the SILEX extraction method (Vilanova et al., 2020). The presence of the transgene was initially verified by detecting red fluorescence derived from the *DsRed* reporter gene using a Leica MZ16 F stereomicroscope (Leica, Wetzlar, Germany). Transgene integration was subsequently confirmed by PCR amplification of a fragment of the Cas12a coding sequence using the primer pair Cas12a_F/Cas12a_R, designed with Primer3 (Table S9). In addition, the genomic regions targeted by the crRNAs in *SmAPRR2* and *SmGLK2* were amplified using their respective primer pairs APRR2_F/APRR2_R and GLK2_F/GLK2_R and subjected to Sanger sequencing to determine the editing status of the regenerated plants and to identify heterozygous mutations. Sanger chromatograms were further analyzed using the Inference of CRISPR Edits (ICE) tool from EditCo (https://ice.editco.bio/#/) to estimate editing outcomes and KO scores in the regenerated lines.

T_0_ plants were self-pollinated to generate T_1_ progeny. To eliminate the transgene, T_1_ seeds lacking *DsRed* fluorescence were selected, as these individuals were expected to be free of the T-DNA insertion. The absence of the T-DNA was also confirmed by PCR using the primer pair Cas12a_F/Cas12a_R (Table S9). Selected seeds were germinated following the protocol described by Ranil et al. (2015). The resulting T_1_ plants were genotyped by Sanger sequencing of the edited loci, and only those carrying homozygous edited alleles were retained for further analysis. This first selection allowed the recovery of transgene-free lines carrying mutations exclusively in *SmGLK2* (*GLK2^cr^*), as well as double-edited lines carrying mutations in both *SmAPRR2* and *SmGLK2* (*APRR2^cr^/GLK2^cr^*). In addition to molecular confirmation, fruit pigmentation was evaluated in the selected lines to verify its consistency with the expected phenotypic effects of the edited loci, namely changes in the green background associated with *SmAPRR2* disruption and changes in the green netting pattern associated with *SmGLK2* disruption. Homozygous lines were subsequently self-pollinated to obtain T_2_ seeds with stable, fixed mutations.

Because of the high editing efficiency observed at *SmGLK2*, no homozygous line carrying an edited *SmAPRR2* allele together with the WT allele at *SmGLK2* (*APRR2^cr^*) was initially recovered among the selected T_1_ progenies, and an additional segregation strategy was implemented to obtain this genotype combination. Selected *APRR2^cr^/GLK2^cr^*lines were crossed with MEL3 WT plants to generate pseudo-F_1_ (pF_1_) progenies, which were heterozygous at the edited loci. These pF_1_ plants were genotyped by Sanger sequencing and subsequently self-pollinated to produce pseudo-F_2_ (pF_2_) populations (Figure S1). A total of 270 pF_2_ individuals were genotyped to identify *APRR2^cr^*lines. Twelve plants with this genotype were selected, and their fruits were phenotypically evaluated to confirm the expected pigmentation pattern.

Finally, six selected homozygous edited lines were retained for subsequent analyses, comprising two *APRR2^cr^* lines, two *GLK2^cr^*lines, and two *APRR2^cr^/GLK2^cr^* lines. The two lines selected within each genotype class carried different edited alleles, thereby capturing allelic diversity within each edited background. Together, these lines represented the three expected fruit color phenotypes: green, netting, and white.

### 4.6. Off-target analysis

Potential off-target sites were evaluated using whole-genome resequencing data from the MEL3 accession as a control and the six selected homozygous edited lines (including a T2 plant derived from T1_3-15, used because DNA of sufficient quality was not available from the original T1 plant). Genomic DNA samples meeting the required quality criteria, with 260/280 and 260/230 absorbance ratios above 1.8, were submitted to BGI Genomics (Hong Kong, China), where 150-bp paired-end libraries were prepared and sequenced on the DNBseq platform. The analysis was performed against the eggplant reference genome “67/3” v3.0 (Barchi et al., 2019), using the six Cas12a crRNAs designed for *SmAPRR2* and *SmGLK2* (Table S10) as query sequences.

First, a genome-wide *in silico* search was carried out with a custom script developed for Cas12a target scanning and executed with Python v. 3.8.3 (Van Rossum and Drake, 2009). The script searched both DNA strands for 23-nt protospacers adjacent to a canonical TTTV PAM and retained candidate sites with up to four mismatches relative to each crRNA. This mismatch threshold was used as a sensitivity analysis to provide a conservative assessment of potential off-target sites (Modrzejewski et al., 2020), while the TTTV PAM was selected because it corresponds to the canonical Cas12a PAM used for crRNA design (Paul and Montoya, 2020). The search window around each off-target candidate was set to ± 1,500 bp to allow subsequent local variant calling in the surrounding genomic region.

Predicted off-target sites were then validated using the resequencing alignments. BAM quality was assessed with Samtools flagstat and Samtools stats v. 1.13 (Li et al., 2009; Danecek et al., 2021). For each predicted off-target site, basewise depth was calculated with samtools depth -a across the protospacer ± 100 bp interval. In parallel, variants were called in the wider ±1,500 bp interval using Freebayes v. 1.3.6 (Garrison and Marth, 2012) with a minimum mapping quality of 30 and a minimum base quality of 20, followed by normalization with BCFtools norm v. 1.3.6 (Danecek et al., 2021). Candidate off-target variants were retained only when they were detected in an edited line, absent in the MEL3 control, supported by a non-reference genotype, and showed an alternative allele count of at least three reads and an alternative allele frequency of at least 0.25. Variants already supported in MEL3 were discarded to avoid interpreting pre-existing MEL3/background polymorphisms as editing-derived changes.

For each predicted off-target site, candidate variants were summarized at the site level, distinguishing all candidate variants absent from MEL3, candidate indels located within the protospacer ± 100 bp interval, and candidate indels overlapping the protospacer. A predicted off-target site was considered sufficiently covered and without detectable evidence of off-target editing when all samples showed at least 95% coverage across the ±100 bp interval and no candidate indel absent from MEL3 was detected in that region.

### 4.7. Flow cytometry analysis

Ploidy level of T_1_ homozygous edited lines was determined by flow cytometry to verify genome stability after the transformation and regeneration processes. For each line, cell nuclei were isolated from three independent young leaves according to the protocol described by García-Fortea et al. (2021). Nuclear DNA content was measured using a CytoFLEX S flow cytometer (Beckman Coulter, CA, USA). The fluorescence intensity of the nuclei was recorded and analyzed using CytExpert to estimate relative DNA content. The WT genotype was included as a diploid reference control, allowing comparison of fluorescence peaks between the edited lines and the WT. Ploidy level was inferred by comparing the position of the main G1 fluorescence peak of each sample, corresponding to diploid nuclei in the G1 phase of the cell cycle, with that of the diploid reference.

## Author contributions

MM-L, AA, JP and PG conceived and designed the study. MM-L, AS and AA performed the experiments. MM-L and AS carried out the molecular and phenotypic analyses. MM-L and PG performed the bioinformatic and statistical analyses. LT, SV, MP, JP and PG contributed resources, protocols, and technical expertise. MM-L prepared the figures and curated the data. MM-L wrote the original draft. All authors reviewed and edited the manuscript. JP and PG supervised the study, managed the project, and acquired funding. All authors read and approved the final version of the manuscript.

## Data availability statement

The data that support the findings of this study are available in the Supplementary files of this article.

