## Supplementary Figures for "CRISPR/Cas12a-mediated allele engineering of *SmAPRR2* and *SmGLK2* reveals complementary control of fruit peel and flesh chlorophyll pigmentation in eggplant"

### Slide 1
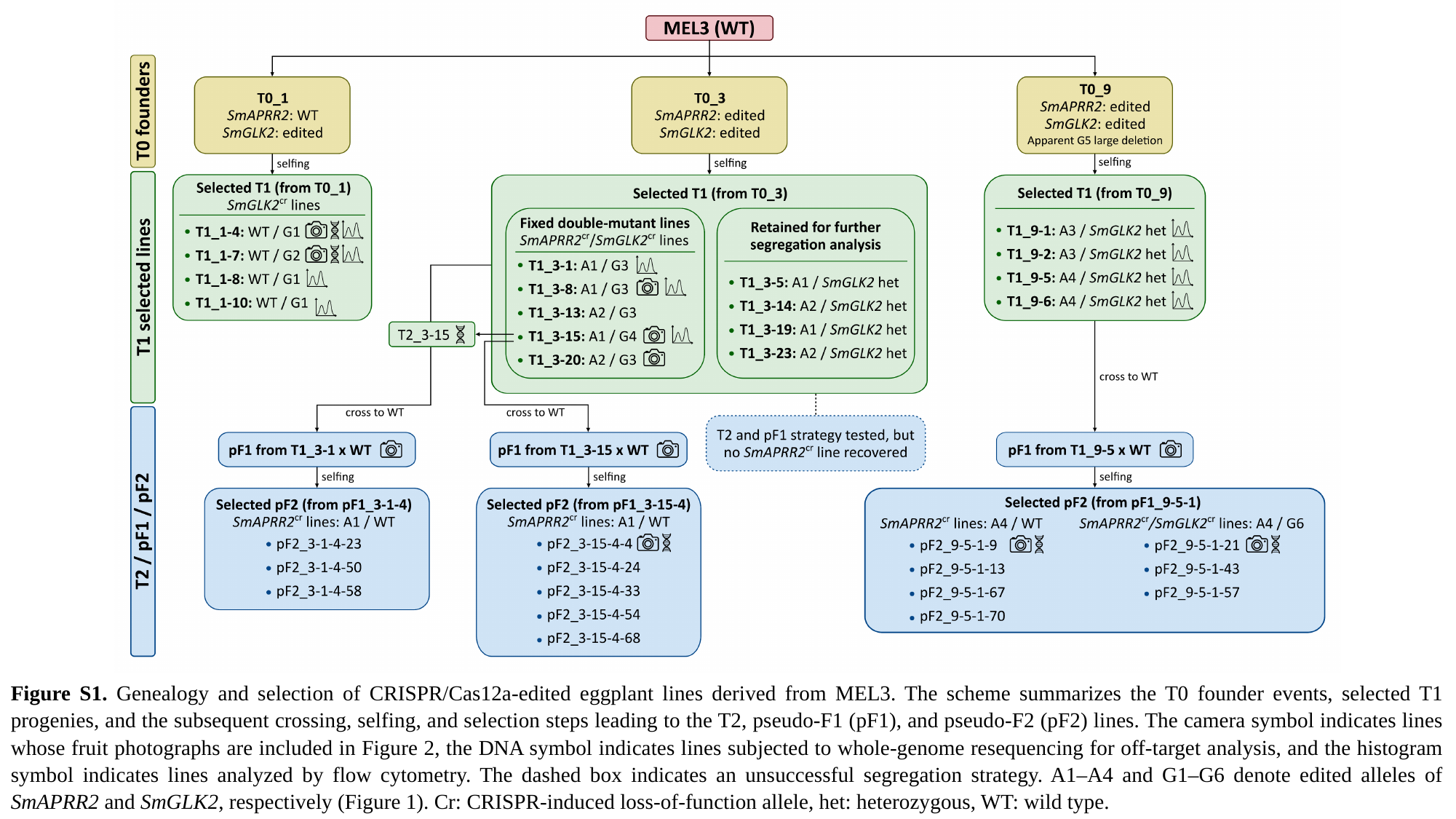

Figure S1. Genealogy and selection of CRISPR/Cas12a-edited eggplant lines derived from MEL3. The scheme summarizes the T0 founder events, selected T1 progenies, and the subsequent crossing, selfing, and selection steps leading to the T2, pseudo-F1 (pF1), and pseudo-F2 (pF2) lines. The camera symbol indicates lines whose fruit photographs are included in Figure 2, the DNA symbol indicates lines subjected to whole-genome resequencing for off-target analysis, and the histogram symbol indicates lines analyzed by flow cytometry. The dashed box indicates an unsuccessful segregation strategy. A1–A4 and G1–G6 denote edited alleles of SmAPRR2 and SmGLK2, respectively (Figure 1). Cr: CRISPR-induced loss-of-function allele, het: heterozygous, WT: wild type.

### Slide 2
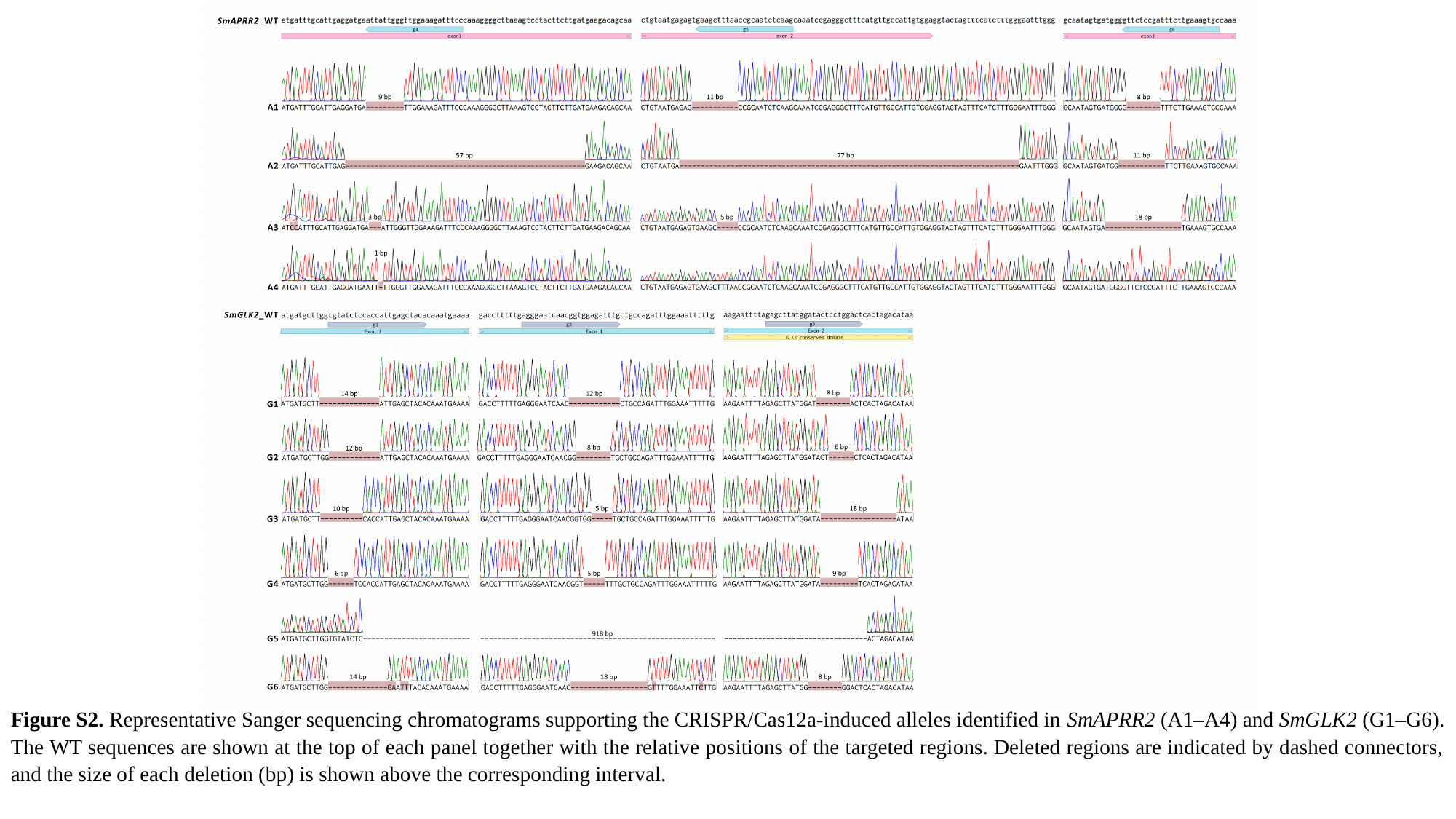

Figure S2. Representative Sanger sequencing chromatograms supporting the CRISPR/Cas12a-induced alleles identified in SmAPRR2 (A1–A4) and SmGLK2 (G1–G6). The WT sequences are shown at the top of each panel together with the relative positions of the targeted regions. Deleted regions are indicated by dashed connectors, and the size of each deletion (bp) is shown above the corresponding interval.

### Slide 3
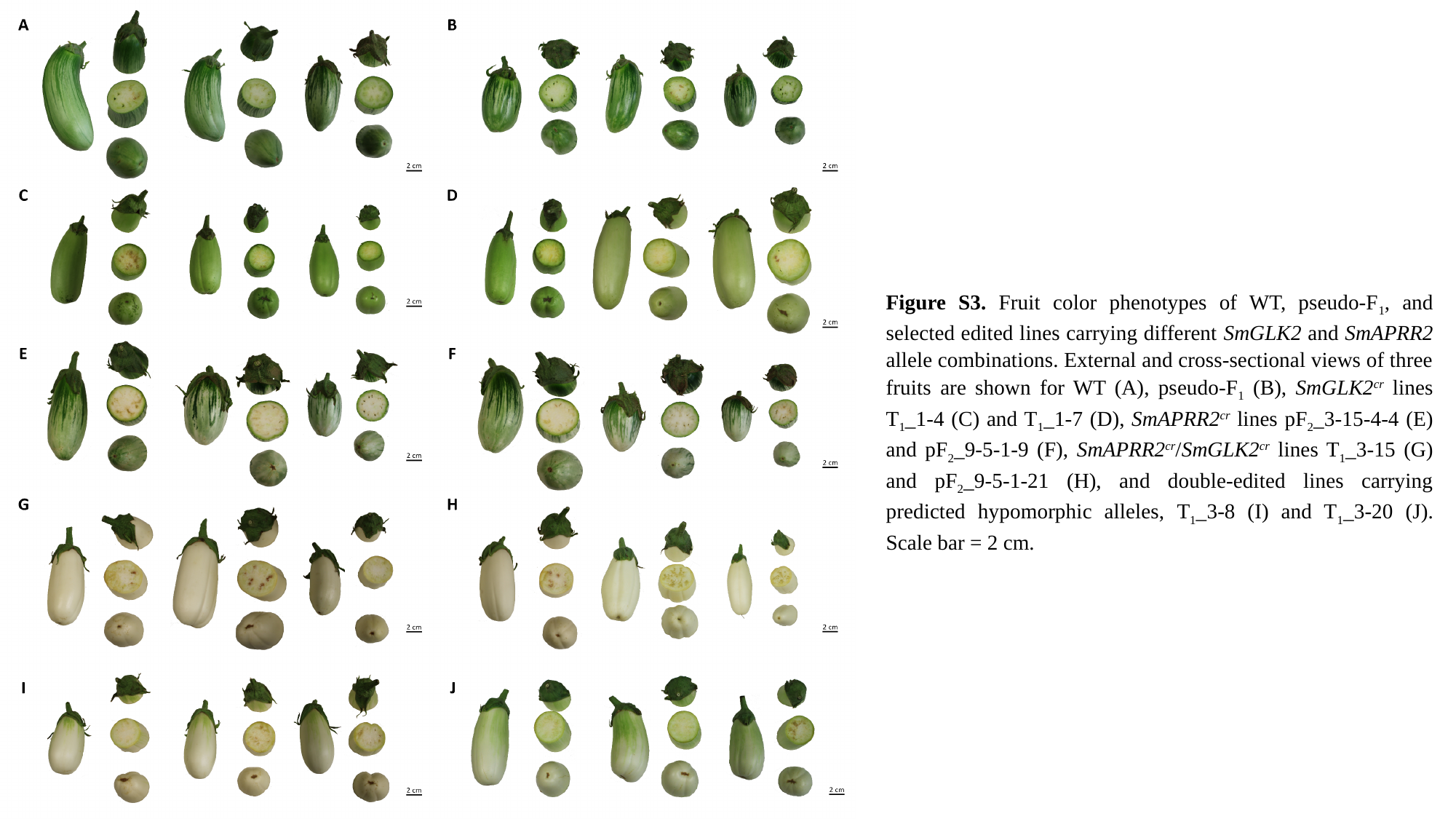

Figure S3. Fruit color phenotypes of WT, pseudo-F_1_, and selected edited lines carrying different SmGLK2 and SmAPRR2 allele combinations. External and cross-sectional views of three fruits are shown for WT (A), pseudo-F_1_ (B), SmGLK2^cr^ lines T_1__1-4 (C) and T_1__1-7 (D), SmAPRR2^cr^ lines pF_2__3-15-4-4 (E) and pF_2__9-5-1-9 (F), SmAPRR2^cr^/SmGLK2^cr^ lines T_1__3-15 (G) and pF_2__9-5-1-21 (H), and double-edited lines carrying predicted hypomorphic alleles, T_1__3-8 (I) and T_1__3-20 (J). Scale bar = 2 cm.

### Slide 4
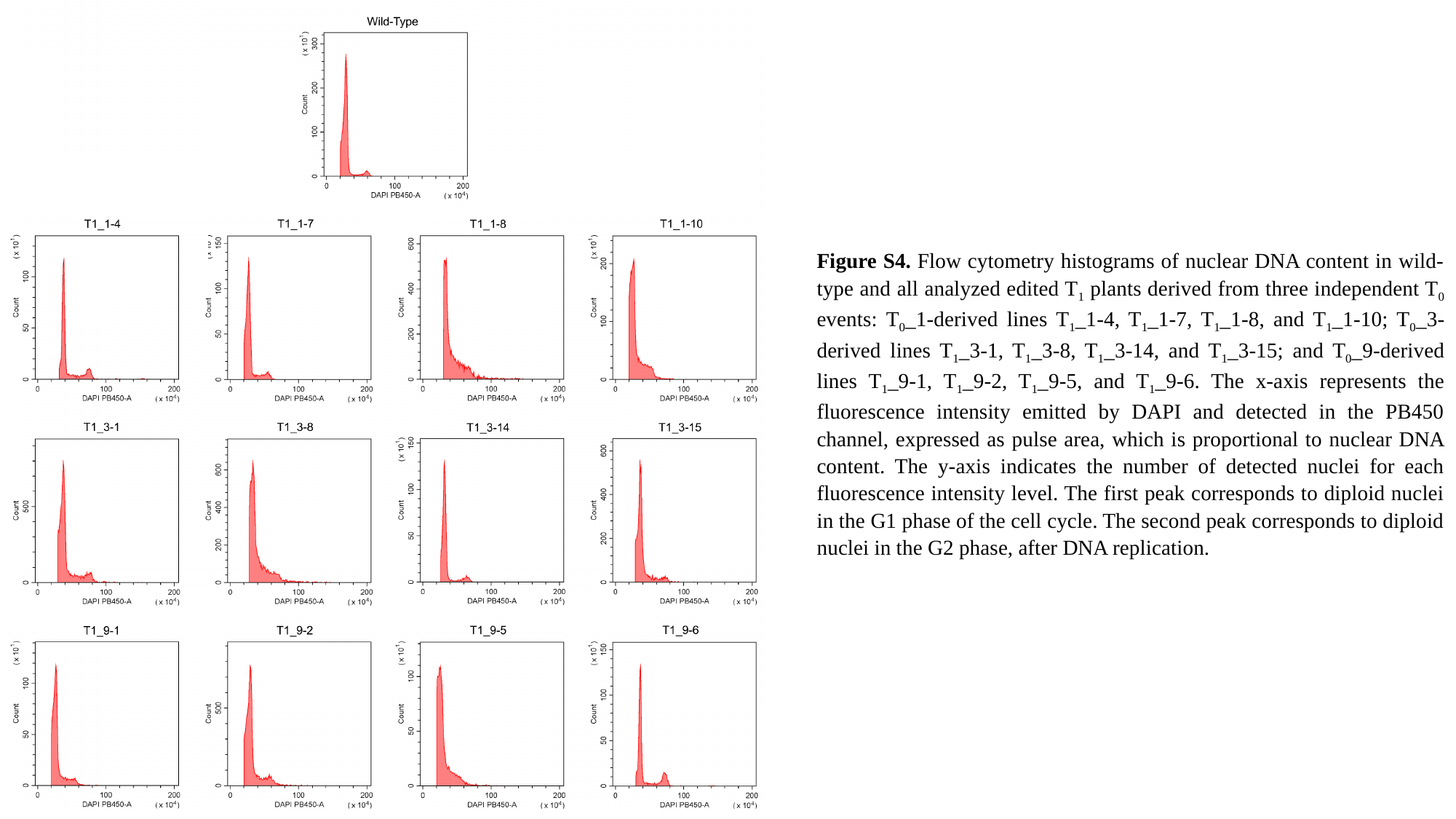

Figure S4. Flow cytometry histograms of nuclear DNA content in wild-type and all analyzed edited T_1_ plants derived from three independent T_0_ events: T_0__1-derived lines T_1__1-4, T_1__1-7, T_1__1-8, and T_1__1-10; T_0__3-derived lines T_1__3-1, T_1__3-8, T_1__3-14, and T_1__3-15; and T_0__9-derived lines T_1__9-1, T_1__9-2, T_1__9-5, and T_1__9-6. The x-axis represents the fluorescence intensity emitted by DAPI and detected in the PB450 channel, expressed as pulse area, which is proportional to nuclear DNA content. The y-axis indicates the number of detected nuclei for each fluorescence intensity level. The first peak corresponds to diploid nuclei in the G1 phase of the cell cycle. The second peak corresponds to diploid nuclei in the G2 phase, after DNA replication.
